# Brown rat demography reveals pre-commensal structure in eastern Asia prior to expansion into Southeast Asia

**DOI:** 10.1101/249862

**Authors:** Emily E. Puckett, Jason Munshi-South

## Abstract

Fossil evidence indicates that the globally-distributed brown rat (*Rattus norvegicus*) originated in northern China and Mongolia. Historical records report the human-mediated invasion of rats into Europe in the 1500s, followed by global spread due to European imperialist activity during the1600s-1800s. We analyzed 14 genomes representing seven previously identified evolutionary clusters and tested alternative demographic models to infer patterns of range expansion, divergence times, and changes in effective population (N_e_) size for this globally important pest species. We observed three range expansions from the ancestral population that produced the Pacific (~4.8kya), eastern China (diverged ~0.55kya), and Southeast (SE) Asia (~0.53kya) lineages. Our model shows a rapid range expansion from SE Asia into the Middle East then continued expansion into central Europe 537 years ago (1478 AD). We observed declining N_e_ within all brown rat lineages from 150-1kya, reflecting population contractions during glacial cycles. N_e_ increased since 1kya in Asian and European, but not Pacific, evolutionary clusters. Our results support the hypothesis that northern Asia was the ancestral range for brown rats. We suggest that southward human migration across China between 800-1550s AD resulted in the introduction of rats to SE Asia, from which they rapidly expanded via existing maritime trade routes. Finally, we discovered that North America was colonized separately on both the Atlantic and Pacific seaboards, yet by evolutionary clusters of vastly different ages and genomic diversity levels. Our results should stimulate discussions among historians and zooarcheologists regarding the relationship between humans and rats.

## INTRODUCTION

The genus *Rattus* originated and diversified in eastern and central Asia, and fossil evidence (Smith and Xie 2008) suggests northern China and Mongolia as the likely ancestral range of the cold-hardy brown rat (*Rattus norvegicus*). Yet, their contemporary distribution includes every continent except Antarctica. As a human commensal, brown rats occupy urban and agricultural areas using food, water, and shelter provided by humans. Rats are one of the most destructive invasive mammals, as they spread zoonotic diseases to humans (Himsworth et al. 2013), damage food supplies and infrastructure (Pimentel et al. 2000), and contribute to the extinction of native wildlife (Harper and Bunbury 2015). As an invasive species, brown rats outcompete native species for resources and are a primary target of eradication efforts (Jones et al. 2016). Brown rats have been domesticated as models for biomedical research with inbreeding leading to disease phenotypes similar to humans (Atanur et al. 2013). Finally, they are a nascent model to study evolution within urban landscapes, as they likely experience multiple selection pressures given their global distribution across a range of habitats and climates (Johnson and Munshi-South 2017).

The historical record indicates that rats colonized Europe in the early 1500s, eastern North America by the 1750s, and the Aleutian Archipelago by the 1780s (Black 1983; Armitage 1993). These historic records provide independent estimates for assessing inferences from demographic models utilizing genomic data. Few other species have archeological or written human records that can be used to corroborate genomic inferences, although house mouse, domestic dogs, and livestock are notable exceptions. Thus, we paired these data sources to test how well demographic models of a rapid and recent global expansion match historic records on rat invasions.

Research into the global expansion of brown rats has focused on both the routes and timings of different invasions; questions of specific interest include the location of the ancestral range, and when rats arrived in Europe. Black rats (*R. rattus*) reached southern Europe by 6kya (Ervynck 2002) and Great Britain by the 300s AD (Yalden 2003), yet brown rats were not recorded in Europe until the 1500s AD. These dates imply vastly different phylogeographic histories for these two commensal rats, likely related to where they speciated within Asia: black rats on the Indian subcontinent and brown rats in the northern steppe. Previous phylogeographic studies of brown rats using mitochondrial DNA identified China as the ancestral range based on private haplotypes and ancestral state reconstructions, with multiple expansions into Southeast (SE) Asia, Europe, and North America (Lack et al. 2013; Song et al. 2014; Puckett et al. 2018). Inference from mitochondria has been limited due to high haplotype diversity observed from locally intense but globally diffuse sampling strategies. Thus, key geographic regions especially around the Indian Ocean basin and the Middle East are unrepresented in current datasets; sampling these areas would allow us to distinguish clinal versus long-distance expansions, where multiple introductions occurred, and mito-nuclear discordance. A phylogeographic analysis using nuclear SNPs inferred hierarchical clustering along five range expansion routes (Puckett et al. 2016). From the putative ancestral range, brown rats expanded southward into SE Asia and eastward into China and Russia (Puckett et al. 2016). The eastward expansion extended to North America with two independent colonizations of the Aleutian Archipelago and sites along the Pacific coast of western North America. From SE Asia rats expanded into Europe (Puckett et al. 2016) via the Middle East (Zeng et al. 2018), where the likely route was aboard ships conducting maritime trade across the Indian Ocean into the Red Sea and Persian Gulf before moving goods onto land. Although these trade routes were established by the 200s BC, they intensified in the 1400-1500s AD (Tucker 2015). The fifth range expansion moved rats to eastern North America, the Caribbean, South America, western Africa, and Australasia during the age of European imperialism of the 1600-1800s (Puckett et al. 2016) with the result that genetic diversity is similar across the Western hemisphere and in western Europe. Ultimately, our previous work inferred the following seven genomic clusters: *Eastern China*, *SE Asia*, *Aleutian*, *Western North America*, *Northern Europe*, *Western Europe*, and (*Western Europe) Expansion*. However, these range expansions were inferred from patterns of population clustering and not specific models that estimate the population tree topology or demographic parameters of the evolutionary lineages. Thus, we generated 10 whole genome sequences (WGS) to represent the previously identified clusters to infer the demographic history of brown rats. We pay particular attention to both divergence times and changes in effective population sizes (N_e_) in relation to climatic changes and human history that may have influenced natural and human-mediated range expansions for this species.

## RESULTS

We sequenced two genomes each from *SE Asia*, *Northern Europe*, *Western Europe*, and the *Western Europe-Expansion* (hereafter-*Expansion*) evolutionary clusters, and one genome each from the *Aleutian* and *Western North America* clusters (NCBI SRA PRJNA344413; Table S1). Average sequencing depth was 28.2× (range 24-38X). We estimated heterozygosity for each individual on the 20 autosomes separately. Samples from *Eastern China* had the highest average chromosomal heterozygosity (0.244) where the *Aleutians* and *Western North America* had the lowest heterozygosity (0.143 and 0.148, respectively; Figure S1).

### Geographic origins of range expansions

We estimated the directionality index (ψ) (Peter and Slatkin 2013) which measures asymmetries between pairwise SFS from 45 global sampling sites genotyped at 32k SNPs to identify the geographic origins and directionality of the different range expansions. We first tested the expansion across Asia and observed that northern sites served as source populations for southward range expansions across the continent (Figure 1A). When we compared SE Asia and the Middle East, we observed that both regions served as source and sink populations, although Z-scores were greater moving from the Middle East to SE Asia (Figure 1B). Given potential connectivity between central Asia and the Middle East, this region requires better sampling to fully describe the regional relationships. The Middle East clearly served as a source of brown rats moving into central Europe, then dispersing across the continent into the Iberian Peninsula, Fennoscandia, and Great Britain (Figure 1C). Since our previous work suggested two expansions into North America, we analyzed the eastern and western seaboards separately. Eastern North America showed a strong signature of expansion from Western Europe (Figure 1D) as expected based on patterns of genomic clustering. Surprisingly, the eastern North America to western North America signatures from genomic clustering analyses (Puckett et al. 2016) were not observed in the directionality index data. Finally, we observed expansion from Russia (i.e. eastern Asia) to both the Aleutian Archipelago and San Diego, USA (*Western North America* cluster; Figure 1E).

**Figure 1.**
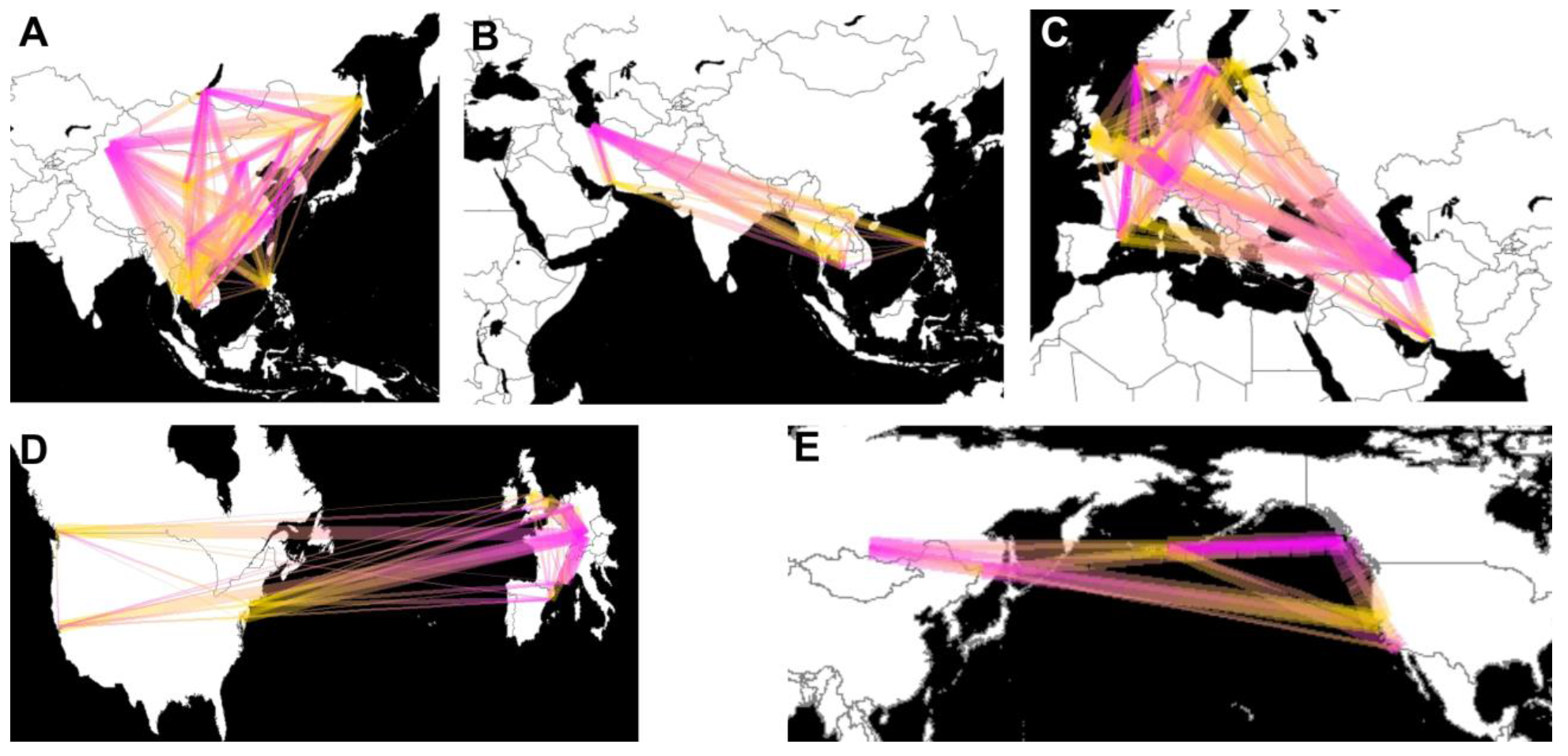
Estimates of regional (A: eastern Asia; B: SE Asia and the Middle East; C: Europe; D: western Europe and the Expansion range, and E: Russia, the Aleutian Archipelago, and Western North America) brown rat range expansions based on pairwise ψ statistics. Lines show directionality from inferred source (pink) to sink (yellow) populations, where thickness was scaled to the Z-score when the absolute value was greater than 5.

### Effective population size through time

We inferred the change in N_e_ over time using the multiple sequentially Markovian coalescent (MSMC) model (Schiffels and Durbin 2014) and scaled the estimates to years and N_e_ using the estimated mutation rate (μ) from the coalescent modeling analysis (see below) of 9.34×10^−8^ and 3 generations per year. (As Deinum *et al.* (2015) estimated μ of 2.96 × 10^−9^ and the precise generation time for rats is unknown, we present alternative estimates of the MSMC model in Figure S2.) We observed two distinct patterns in the MSMC results related to the Pacific (*Aleutian* and *Western North America*) and all other clusters. The Pacific clusters declined sharply in N_e_ beginning approximately 50kya (Figure 2). MSMC is not accurate in its last two time periods, therefore we present N_e_ of the third time lag which was approximately 200 years ago and estimated at 1,460 and 1,550 effective individuals respectively in the *Aleutian* and *Western North America* clusters (Figure 2).

**Figure 2.**
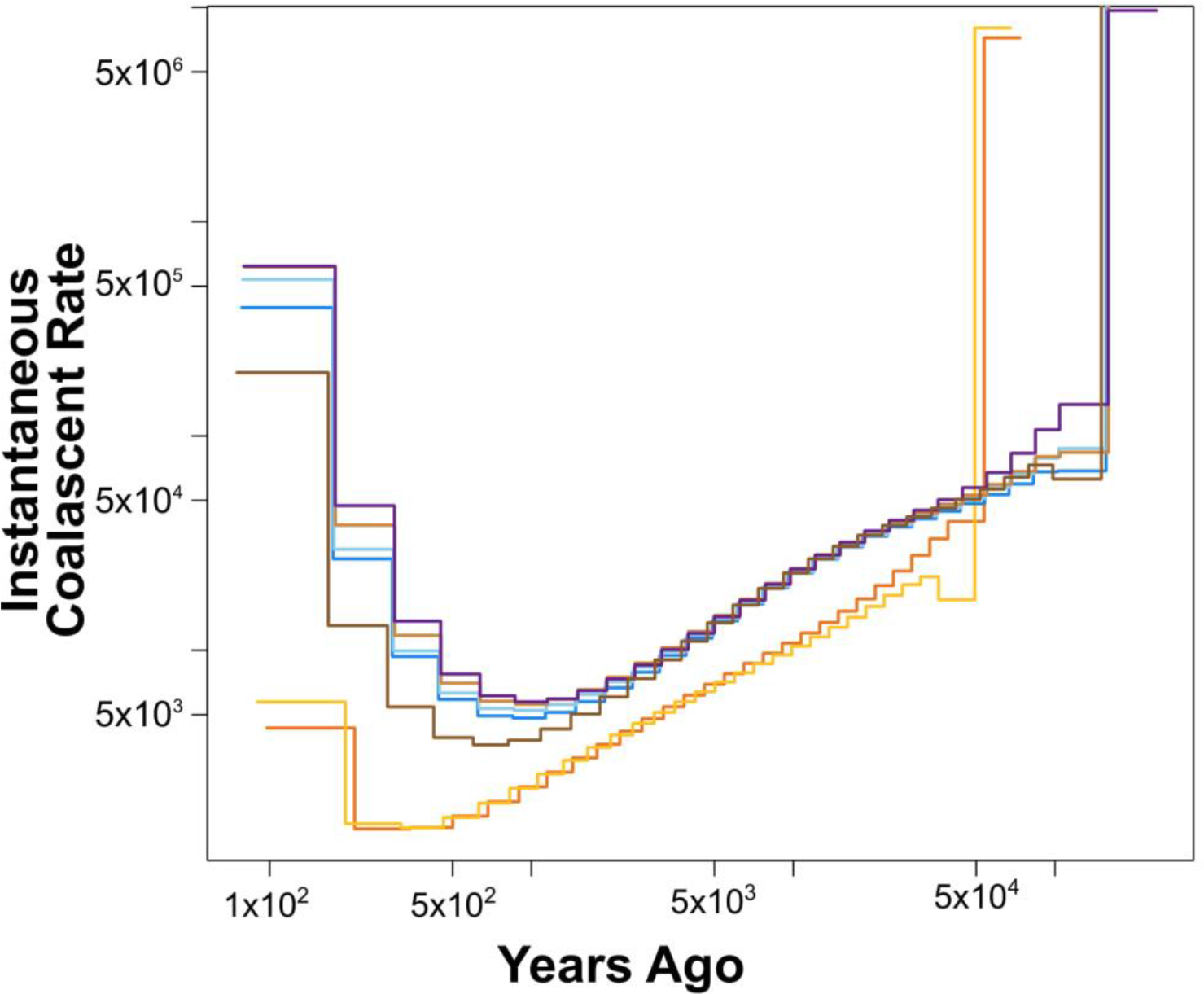
Plot of change in the instantaneous coalescent rate over time using MSMC where the x-axis is years before the present. Each evolutionary cluster was represented by a different color: *eastern China*- dark brown; *SE Asia*- light brown; *Aleutian*- orange; *Western North America*- yellow; *Northern Europe*- purple; *Western Europe*- light blue; and *Expansion*- medium blue.

The second pattern was concordant between *Eastern China*, *SE Asia*, *Northern Europe*, *Western Europe*, and *Expansion* clusters. N_e_ steadily declined from approximately 150 - 1kya before increasing in the most recent time periods (Figure 2). Approximately 200 years ago (the first reliable time step), N_e_ was: 13,000 in *Eastern China*, 38,000 in *SE Asia*, 47,000 in *Northern Europe*, 29,000 in *Western Europe*, and 26,000 in *Expansion* (Figure 2).

### Demographic model

Based on previous work on the hierarchical genetic clustering of brown rats (Puckett et al. 2016) and the range expansion results, we split the range into Asian and European derived clusters and inferred that SE Asia linked the two regions. Thus, we built our full demographic model by conducting model selection in two stages where we first identified the models that best represented divergence patterns in Asia (Figure S3) and Europe (Figure S4) separately, then combined those tree topologies into a global model for parameter estimation. The best Asian model had an ancestral unsampled population with independent divergence events for *Eastern China, SE Asia*, and the “Pacific” cluster that diverged into the *Aleutians* and *Western North America* (Figure S3). For the European model, the best supported model used *SE Asia* as the ancestral population then inferred a series of divergences first into the Middle East then *Western Europe*, followed by independent divergences of *Northern Europe* and the *Expansion* from *Western Europe* (Figure S4).

Using WGS data from 14 genomes, we modeled the nine-population topology inferred from the sub-models (Figure 3). We ran five models varying the rate parameters that allowed growth or contraction of N_e_ over time given our MSMC estimates (Figure 2), and observed that the best-supported model included decreasing ancestral population size and increasing size since the start of the range expansions (Appendix 1). We estimated that *Eastern China* diverged from the ancestral population 554 years ago (90% highest density probably [HPD]: 513 - 8666 years; Table 1). The Pacific cluster diverged from the ancestral population 4.8kya (HPD: 3.1 - 12.3kya), then the *Aleutians* and *Western North America* diverged 2.6kya (HPD: 1.0 - 9.1kya). The divergence that led to the global expansion of rats occurred rapidly, where rats first expanded into *SE Asia* 538 years ago (1477 AD; HPD: 1475-1745 AD). Our model also estimated 538 years (HPD: 1471-1741 AD) for the entry of rats the Middle East. We estimated rapid divergence of rats into Europe including the *Western Europe* divergence 537 years ago (1478 AD; HPD: 1472-1741 AD), and *Northern Europe* divergence 536 years ago (1479 AD; HPD: 1470-1739 AD). Finally, we estimated the *Expansion* cluster diverged 509 years ago (1494 AD; HPD: 1487-1741 AD).

**Figure 3.**
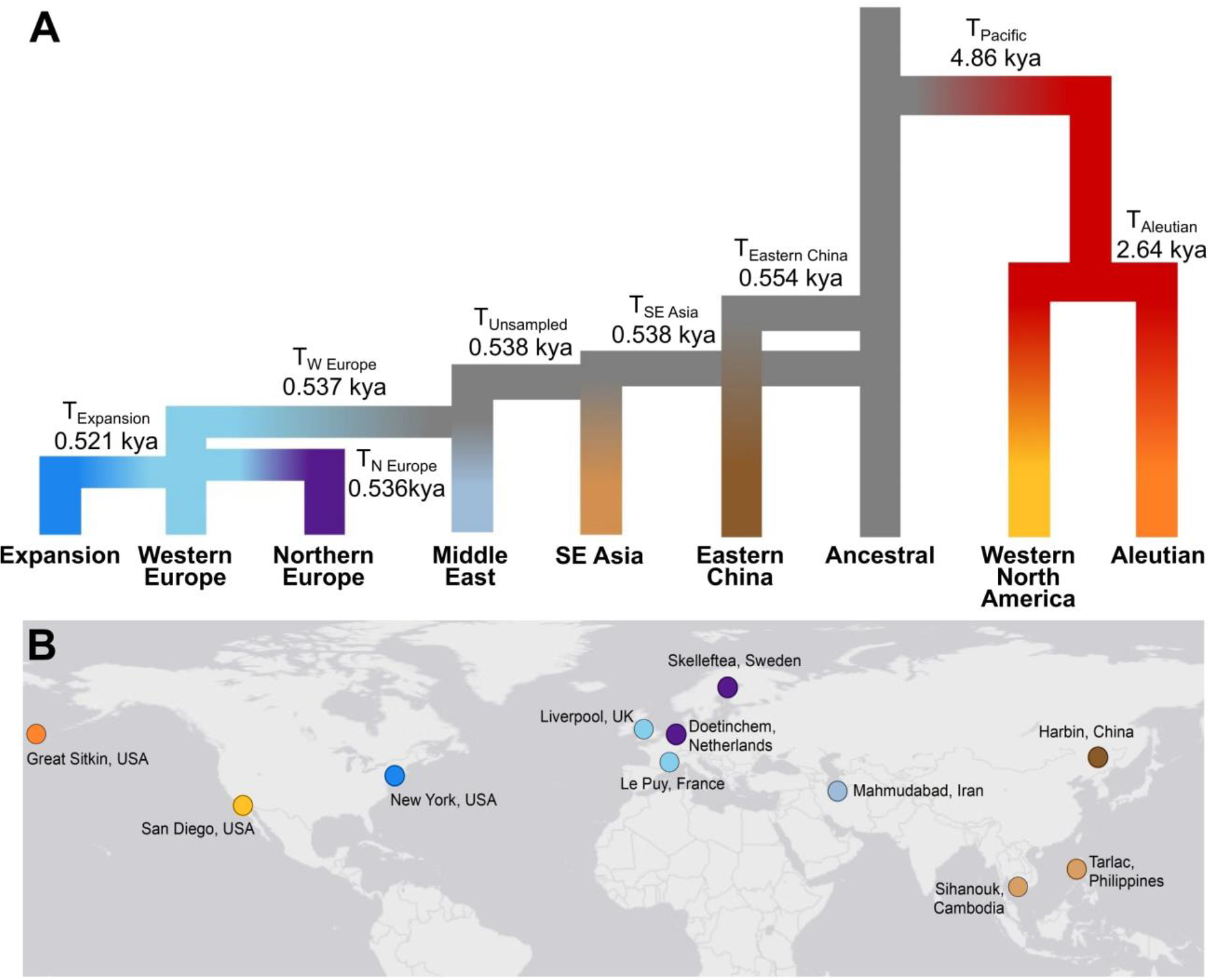
(A) The best supported demographic model contained nine evolutionary clusters inclusive of two unsampled populations. The divergence times in generations and N_e_ are listed in Table S4. (B) Map of global sampling locations for the WGS demographic model where evolutionary clusters were represented by different colors: *Eastern China*- dark brown; *SE* Asia- light brown; *Aleutian*- orange; *Western North America*- yellow; Middle East- grey-blue; *Northern Europe*- purple; *Western Europe*- light blue; and *Expansion*- medium blue.

**Table 1.** Parameter estimates from the best supported model of global brown rat demography for eight evolutionary clusters and an unsampled ancestral populations using 14 whole genomes. Both point and 90% highest density probability (HPD) estimates are presented for each model parameter (N- population size, R- rate of population change, T- divergence time, and μ- mutation rate). See Appendixes 1 and 2 for model specifications and Figure 2A for population topology.

| Parameter | Units | Estimate | 90% HPD |
| --- | --- | --- | --- |
| N<sub>Ancestral</sub> | 2N | 129714 | 43297 - 231318 |
| N<sub>Eastern China</sub> | 2N | 2096 | 665 - 45667 |
| N<sub>Aleutians</sub> | 2N | 34627 | 8955 - 40705 |
| N<sub>Western North America</sub> | 2N | 22366 | 5356 - 53122 |
| N<sub>SE Asia</sub> | 2N | 1547 | 448 - 1702 |
| N<sub>Iran</sub> | 2N | 268 | 158 - 635 |
| N<sub>Western Europe</sub> | 2N | 854 | 267 - 1314 |
| N<sub>Northern Europe</sub> | 2N | 836 | 259 - 1204 |
| N<sub>Western Europe Expansion</sub> | 2N | 423 | 279 - 1019 |
| R<sub>Ancestral</sub> | | $7.54 \times 10^{-2}$ | $0.07 - 1.68 \times 10^{-3}$ |
| R<sub>Pacific</sub> | | $2.53 \times 10^{-4}$ | $0.19 - 3.97 \times 10^{-4}$ |
| R<sub>Ghost</sub> | | $-1.84 \times 10^{-5}$ | $-9.17 - -0.03 \times 10^{-5}$ |
| R<sub>Eastern China</sub> | | $1.07 \times 10^{-4}$ | $-3.85 - -0.83 \times 10^{-3}$ |
| R<sub>Aleutians</sub> | | $-9.14 \times 10^{-4}$ | $-5.58 - -7.13 \times 10^{-4}$ |
| T<sub>Eastern China</sub> | gen | 1663 | 1541 - 25999 |
| T<sub>Pacific</sub> | gen | 14580 | 9330 - 36987 |
| T<sub>Aleutian - Western North America</sub> | gen | 7926 | 3107 - 27317 |
| T<sub>SE Asia</sub> | gen | 1615 | 809 - 1618 |
| T<sub>Iran</sub> | gen | 1613 | 822 - 1630 |
| T<sub>Western Europe</sub> | gen | 1612 | 821 - 1629 |
| T<sub>Northern Europe</sub> | gen | 1608 | 828 - 1634 |
| T<sub>Western Europe Expansion</sub> | gen | 1563 | 823 - 1584 |
| $\mu$ | mutations gen<sup>-1</sup> | $9.34 \times 10^{-8}$ | $8.15 - 9.89 \times 10^{-8}$ |

We ran the cross-coalescence analysis within MSMC2 to estimate the rate of divergence between the seven clusters with high depth of coverage (i.e. excluding Iran). We observed that divergence was complete between both the *Aleutians* or *Western North America* and all other populations (Figure S5). The European clusters showed similar patterns of divergence with *Eastern China* with approximately 60% divergence complete (Figure S5A). Cross-coalescence between the *Aleutians* and *Western North America* increased approximately 200 generations ago before decreasing to 50% (Figure S5C). The four clusters making up the most recent expansions (*SE Asia*, *Northern Europe*, *Western Europe*, and *Expansion*) had signatures of increasing cross-coalescence over the past 1,000 generations (Figure S5 B, D-F). We believe that the rapid global range expansion resulted in high rates of coalescence events between instead of within evolutionary clusters that produced this signature in the analysis.

### ddRAD Demographic Models

We used a ddRAD-Seq dataset (Table S2) to further investigate regional population tree topologies to better understand patterns of global range expansion; below we detail the motivation and results for each analysis. We estimated divergence time of the two Pacific clusters at 2.6kya (Table 1) which was surprisingly old given the historic record that rats were introduced to the Aleutian Archipelago in 1780 AD. Thus, we estimated the population tree topology between eastern Asia and western North America (Figure S6). We observed that a model where eastern China and Russia were sister populations with an admixture pulse from Russia into Adak Island (*Aleutian* cluster) of 30% occurred 215 generations ago (1943 AD) was the best supported model. The estimated timing of this admixture pulse was younger than the historic record predicted.

Our previous clustering results suggested that brown rats in the Philippines were diverged from other SE Asia countries, and that there may be gene flow between Thailand and Cambodia (Puckett et al. 2016); therefore, we modeled the population tree within SE Asia. We observed that the Philippines were well diverged from mainland populations, and that gene flow from Thailand into Cambodia was present (Figure S7). The population tree topology supported the geography where Cambodia and Vietnam were sister populations that shared an ancestor with Thailand (Figure S7).

We split European populations between the *Western* and *Northern* evolutionary clusters and observed patterns concordant with geography; specifically, Norway and Sweden were sister populations and shared a common ancestor with the Netherlands on continental Europe (Figure S8). Similarly, France and Spain on the Iberian Peninsula shared a common ancestor with Great Britain, an island nation (Figure S9).

North America presents the most complex scenario as invasion occurred on both the east and west coasts, and shows patterns of cross-continent range expansions in both directions (Figure 1D-E) (Puckett et al. 2016). We modeled the population tree of North America using the following populations: Udon Thani, Thailand (*SE Asia*), Nottingham, Great Britain (*Western Europe*), NYC, USA (*Expansion*), San Diego, USA (*Western North America*), and ran each topology independently adding in either Vancouver, Canada (*Expansion*) or Berkeley, USA (*Western North America*) to understand variation along the Pacific seaboard. Our previous work (Puckett et al. 2016) identified that brown rats in Vancouver had high proportions of European ancestry with some Asian ancestry; we interpreted this result as original invasion by the *Expansion* cluster with gene flow from neighboring Pacific coast populations that contained *Aleutian* or *Western North America* ancestry. Our best supported model showed admixture between the *Expansion* and *Western North America* clusters (Figure S10); surprisingly, the proportion from the *Expansion* cluster was 36% which was low compared to our previous result of ~90% European ancestry. The pattern in Berkeley, USA differed from that in Vancouver, where a model of population divergence between San Diego (*Western North America*) and Berkeley was observed prior to an admixture pulse from NYC (*Expansion*; Figure S10). This admixture pulse was estimated as 9% of the total Berkeley ancestry which was surprising given previous estimates of high proportions of European ancestry.

## DISCUSSION

Our demographic modeling inferred that brown rats expanded from an ancestral range in northern Asia into eastern China, western North America, and SE Asia (Figures 1,2, and S1). We included an unsampled ghost population in our model to represent this ancestral range in northern Asia. Brown rat fossils have been described from northern China and Mongolia (Smith and Xie 2008), and our range expansion results (Figure 1A) suggest eastern Russia as a possible part of the ancestral range. The Pacific cluster diverged earliest from the ancestral population 4.8kya, and divergence of the *Aleutian* and *Western North America* clusters occurred 2.6kya (Figure 3, Table 1). Our cross-coalescence analysis (Figure S5C) suggested that divergence between these clusters may be as recent as 100 generations ago, which would explain why the patterns of change in N_e_ were similar over time. These results also suggest an explanation for the wide HPD estimates for these clusters in our demographic model (Table 1). We emphasize that the divergence of these clusters does not identify the timing of the introduction to the Aleutian Archipelago or the Pacific coast of North America where samples were collected. The historic record indicates rats were moved to the Aleutian Archipelago by Russian fur traders in the 1780s (Black 1983). Our regional population model suggested a scenario with gene flow from a population in eastern Russia into Adak Island (Figure S6), thereby suggesting two introductions of rats onto Adak. Our results suggest that more work is needed to understand invasion patterns of rats into western North America from eastern Asia. Specifically by incorporating both more sites and ancient samples, demographic models could estimate divergence times to differentiate between models of contemporary or historic movement of rats; this question is particularly interesting because human migration into the region occurred approximately 36kya (Moreno-Mayar et al. 2018). Thus, when rats first moved across Beringia remains an open question.

We estimated that the expansion across Asia was rapid and recent. Our modeling supported an independent expansion into *Eastern China* from the ancestral range 554 years ago (Figure 3A, Table 1). Given the eastern location of Harbin, China (where this lineage was sampled), we felt that it was reasonable to assume this was an independent expansion instead of part of the broader southern expansion into SE Asia (Figures 1A and 3A). Interestingly, the Harbin population contains high mitochondrial diversity, with the most divergent clades estimated to 96kya (HPD: 70-128kya) (Puckett et al. 2018). This high mitochondrial diversity may reflect movement from multiple ancestral populations into the eastern portion of the range prior to recombination creating a unique nuclear genomic signature for the lineage.

We estimated that the *SE Asia* cluster diverged from the ancestral population 0.538kya (1477 AD, Figure 3, Table 1). The timing of this divergence immediately raises the question of why rats did not expand sooner, as overland trade between China and SE Asia was established by the 500s AD (Lieberman 2009); maritime trade between these regions and the Indian Ocean basin was established before the 900s AD (Heng 2009). A partial explanation may be due to the intersection of climate and human demography across eastern Asia. The Medieval Climate Anomaly (850-1250 AD) aided agricultural expansion and human demographic growth in China, specifically prompting urban centers to expand outward at a time of human movement from northern arid lands to more agriculturally productive lands in the south (see references within Lieberman 2009). However, the end of this climatic period resulted in drought, famine, political instability, and ultimately human demographic contractions in both China and SE Asia; fortunes reversed in the late 1400s to mid-1500s as the climate improved and populations expanded again (Lieberman 2009). We hypothesize that this southward human demographic expansion facilitated the range expansion of brown rats, which explains the clinal pattern of ancestry from northern China across southern China into SE Asia that Zeng *et al.* (2018) observed. Thus, the founding of new agrarian communities and increasing inter-connectedness with urban centers would serve as stepping-stones for rats to move from northern China to SE Asia during the two periods of human demographic expansion.

Our results regarding an ancestral range in the north with southward expansion into SE Asia stand in marked contrast to a different study that identified brown rats in SE Asia as the ancestral population with a northward expansion (Zeng et al. 2018). Both analyses used coalescent modeling approaches but with four primary differences: independent datasets, mutation rates, generation time, and tree topology. To address variation in the mutation rate, we ran our global model with the mutation rate fixed to 1.103 × 10^−9^ as used by Zeng et al. (2018) and observed a decreased model fit than when we allowed the mutation rate to be estimated as part of the model; therefore, we reported the model using the estimated rate of 9.34 × 10^−8^. Regarding the generation time, we converted generations to years using the estimate of 3 generations per year where Zeng et al. (2018) used 2 generations per year. Thus, if both papers estimated a divergence given the same number of generations, the estimate of 3 generations per year would make those estimates more recent in time and 2 generations per year would estimate the event further back in time. Without direct field observations of rat fecundity and how it may vary with resources or climate, we are unable to identify the exact dates and thus acknowledge that discrepancy between the papers.

While the factors above likely contributed to small differences between our results and that of Zeng et al. (2018), we think that the more substantial discrepancy results from the population tree topology. Specifically, Zeng and colleagues (2018) included admixed samples containing SE Asian ancestry (that were geographically located in southern China) within the northern China cluster. By grouping the mixed ancestry samples preferentially with the northern Asia cluster, the coalescent model was better supported when the SE Asia population was ancestral. This result was likely due to better model fit given that a small proportion of SE Asian alleles were within the northern Asia cluster. We believe their result was not due to the true history of the populations but can instead be attributed to artefacts introduced by inappropriate sample clustering. Finally, we observed that inclusion of an unsampled ancestral population improved our model fit. Unsampled populations can influence parameter estimates of N_e_ and migration rates and has been shown to improve or at least not harm parameter estimation within the full model (Beerli 2004). Adding an unsampled population to our model was important given the limited number of chromosomes genotyped, as large sample sizes decrease the effect of unsampled populations on parameter estimates (Slatkin 2005).

We estimated a rapid range expansion from SE Asia into Europe via the Middle East (Table 1). There was concordance between the phylogeographic patterns in our results and those of Zeng and colleagues (2018); however, we estimated the divergence time into the Middle East 538 years ago (Table 1) and they 3,100 years ago (2,066 years when using 3 generations per year). These discrepancies were likely due to how population size and divergence time interact in coalescent models; specifically, we observed greatly improved model fit when including an ancestral population rate change parameter (R_Ancestral_; Appendix 1). It was unsurprising that our model also estimated significantly smaller lineage specific N_e_. Our estimate of N_e_ in NYC, USA (211 individuals; *Expansion* cluster; Table 1) was similar to an independent analysis of rats across NYC that estimated N_e_ of 260 individuals (Combs et al. 2018); therefore, we have high confidence in our model estimates of N_e_, and believe that the pattern of population size change from the MSMC analysis was informative even if the exact estimates were high. Finally, we estimated that the *Expansion* cluster diverged from *Western Europe* around 1494 AD (Table 1). This estimate was much older than that of the historic record (Armitage 1993) and either indicates limits to parameter estimation for recent divergence events or an area for improvement within our model.

### Ancestral population size

We observed that N_e_ steadily declined in both the Pacific and Ancestral range populations around 150kya and 50kya, respectively (Figure 2). These declines began prior to the Last Glacial Maximum (22 - 18kya), a climatic period when populations of many species declined due to range contractions and/or shifts. More recent increasing population size appears related to the demographic and geographic range expansion mediated by rats commensal relationship with humans instead of climatic events alone.

### Range expansion via human-mediated movements

Our results identified that the global range expansion of rats occurred recently (1470s AD) and rapidly from SE Asia into Europe via the Middle East and was likely linked by maritime trade between those regions. This stands in marked contrast to previous assumptions that brown rats were transported westward along the Silk Road through central Asia into Europe. This is counterintuitive as overland trade routes from central China to Persia were established 2.1kya (105 BC) and goods reached Rome by 46 BC (Tucker 2015). The Silk Road passed through part of the native range of brown rats, unlike black rats that originated on the Indian subcontinent (Aplin et al. 2011). Assuming that rats evolved their commensal relationship with humans prior to their global range expansion, as observed with house mouse (Suzuki et al. 2013), the availability of cities, road networks, and a flow of merchants naturally suggests a way to expand westward. The Silk Road may have not been the route for expansion due to the limited distance that merchants traveled along the route, as goods went further than the caravans containing the resources rats would need for survival (Tucker 2015). Further, high aridity and lack of water sources may have limited rat movement via the Silk Road. Yet this does not preclude the idea that brown rats may have expanded westward via Silk Road cities and were then extirpated due to the collapse of those cities during changing geo-politics and shifts towards maritime trade (Tucker 2015). We instead suggest that pulses of southward human demographic expansion from northern China during favorable climatic conditions enabled the expansion of rats into SE Asia from which they expanded westward. This hypothesis was supported by our range expansion models (Figure 1) showing westward movement from the Middle East into central Europe, then expansion in all directions across Europe. We present this historical narrative as a hypothesis supported by our demographic model, but also to stimulate interest in further study by historians and zooarchaeologists to examine the historical expansion of this globally important invader.

## MATERIALS and METHODS

### Whole genome sequencing and datasets

We selected 10 individuals for whole genome sequencing: two each representing evolutionary clusters within *SE Asia* (Philippines and Cambodia), *Northern Europe* (Sweden and Netherlands), *Western Europe* (England and France), and *Expansion* (New York, USA), and one sample each from the *Aleutian Islands* and *Western North America* (Table S1, Figure 2B). We generated paired-end reads for each sample (4ng RNase treated genomic DNA) by sequencing on an Illumina HiSeq 2500 at the New York Genome Center. Initial bioinformatics were completed by the New York Genome Center where genomes were mapped to the Rnor_5.0.75 reference (Gibbs et al. 2004) using BWA-MEM v0.7.8 (Li and Durbin 2010). Then, duplicates were marked using Picard Tools v1.122 and indels were realigned with the GATK v3.4.0 IndelRealigner (McKenna et al. 2010). We sorted and indexed BAM files using SAMTOOLS v1.3.1 (Li et al. 2009). Data for these 10 genomes are available on the NCBI SRA BioProject PRJNA344413 (Puckett et al. 2018).

We combined these 10 new WGS sequences with three existing datasets (Table S1) depending on the analysis. Specifically, we downloaded whole genomes from 11 brown rats and one black rat (*R. rattus*) collected in Harbin, China (ENA ERP001276), although to not bias estimates with unequal sample sizes we ran analyses using only Rnor13 and Rnor14 (Deinum et al. 2015) which were randomly selected. We downloaded 54 low-depth WGS brown rats collected in cities across Russia, China, and Iran (Beijing Institute of Genomics BioProject CRA000345, accessions: CRR021172 - CRR021339) (Zeng et al. 2018). Two of these samples (Iran5 and Iran9) were used in WGS analyses, whereby we mapped the raw reads to the Rnor_5 reference with Bowtie v2 (Langmead and Salzberg 2012) using the default parameters, then sorted and indexed in SAMTOOLS. All 54 genomes were mapped to the Rnor_6 reference with Bowtie v2, sorted and indexed using SAMTOOLS, then had a set of 32k SNPs extracted using a position list in SAMTOOLS to make the data comparable to genotypes from 326 brown rats collected from around the globe (Puckett et al. 2018). Using these data sources, we created four datasets which varied in input samples and processing depending on the resultant analysis; we describe the input data and analyses in detail below.

### Patterns of Range Expansion

We explored the geographic patterns of the global range expansion using the directionality index (Peter and Slatkin 2013) calculated from the site frequency spectra (SFS). The directionality index identifies the expected geographic location that acted as the center of a range expansion event. Where alternative tree topologies may be tested with demographic models to identify the one with the highest likelihood to the observed data.

This analysis utilized the combined ddRAD-Seq genotypes from Puckett et al (2018) and WGS data from Zeng et al. (2018) at 32k SNPs. We removed sampling sites represented by a single individual for a final data set containing 276 individuals from 45 locations. The VCF was converted into PLINK format, then imported into the *rangeExpansion* package for R (Peter and Slatkin 2013). We calculated the directionality index, ψ, for all population pairs using the get.all.psi function. To determine significance, we calculated the standard error of the upper triangle of the pairwise ψ matrix excluding the diagonal, thereby allowing us to calculate the Z-score for each population. For each region of interest, we plotted data for each pair of populations where the absolute Z-score was greater than 5 and visually assessed the geographic patterns of source and sink populations.

### Estimates of N_e_ Through Time

We estimated the change in effective population size over time in each evolutionary cluster using MSMC2 (Schiffels and Durbin 2014). To call variants, we used SAMTOOLS mpileup across all samples (10 WGS genomes sequenced here and two Chinese genomes) with a minimum mapping quality of 18 and the coefficient to downgrade mapping qualities for excessive mismatches at 50. We then utilized the variant calling in BCFTOOLS v1.3 with the consensus caller and excluded indels which limited the dataset to bi-allelic SNPs, before pipping the output to the authors’ bamCaller.py script that produced per chromosome masks and VCF files for each individual. As there was not a brown rat reference panel, we phased the 12 individuals plus two inbred lines (SS/Jr and WKY/NHsd; NCBI SRA accessions ERR224465 and ERR224470, respectively (Atanur et al. 2013)) for each of the 20 autosomes using fastPHASE v1.4.8 (Scheet and Stephens 2006). We generated genome-wide masks for each chromosome using SNPable (Li 2009), then converted to a bed file with the makeMappabilityMask.py script. Finally, we used the generate_multihetsep.py script to create the MSMC2 input files before running the program within and between population clusters. Specifically, we estimated change in N_e_ over time for each of the seven evolutionary clusters using two haplotypes for the *Aleutian* and *Western North American* clusters and four haplotypes for each other cluster. We also estimated the proportion of population divergence over time using the cross-population analysis, and combined results from individual populations with the cross-population analysis using the combineCrossCoal.py script provided.

### WGS Demographic Modeling

We inferred the demographic history of rats by modeling alternative scenarios that compared the observed and expected site frequency spectra (SFS) for each evolutionary cluster. We combined the 10 genomes sequenced in this study, two genomes from Harbin, China, and two genomes from Mahmudabad, Iran (Table S1). We limited SNP calling to sites observed in 10 of 12 genomes (-minInd; excluding those from Iran which had lower depth of coverage), to the 20 autosomes, and to bases that had a minimum mapping quality (-minmapq) of 30 and minimum Q score (-minQ) of 20 using ANGSD v0.915 (Korneliussen et al. 2014). We estimated genotype likelihoods using the function implemented in SAMTOOLS (-GL 1) (Li et al. 2009). The *R. rattus* individual from China served as the outgroup allowing for identification of ancestral and derived alleles; then we calculated a folded SFS for each pairwise evolutionary cluster in ANGSD.

Given the large number of evolutionary clusters to model, we first modeled the relationship between *Eastern China*, *SE Asia*, *Aleutian*, and *Western North America* by comparing five four-population models and five five-population models that included an unsampled population (Figure S3; Appendix 2). The best supported scenario (Model 6 in Figure S3) had a topology that included an ancestral unsampled ghost population with independent divergence of *Eastern China*, *SE Asia*, and the Pacific clusters. We then modeled four populations of a five-tree topology between the Middle East, *SE Asia*, *Northern Europe*, *Western Europe*, and the *Expansion*. Our previous work on brown rat phylogeography suggested that rats expanded into Europe from SE Asia (Puckett et al. 2016) and Zeng et al (2018) showed that the Middle East served as an intermediary point between SE Asia and Europe; thus, we tested the topology between the three European clusters (Appendix 2). The best supported scenario (Figure S4) had an initial divergence of *Western Europe* from the Middle East, with *Northern Europe* and the *Expansion* diverging independently. For initial testing models, we did not allow population size to change through time, and we set the mutation rate at 2.5 × 10^−8^ mutations per generation.

The best supported sub-models were concordant with the range expansion results; thus, we combined the topologies into a nine-population model. We tested this global model with no within lineage change in N_e_, as well as allowing N_e_ to vary for both tip and ancestral branches (Appendix 1). Unlike in the sub-models described above, we estimated the mutation rate parameter within the model. We used three generations per year to convert parameter estimates; all time calculations were done since 2015.

We ran 50 iterations of the nine-population model in fastsimcoal2 v2.6.0.3, then identified the iteration with the highest estimated likelihood. Using these point estimates, we generated 500 samples of pairwise SFS each containing 50,000 markers that served as pseudo-observed data for estimating parameter ranges under the best supported model. We calculated the 90% highest probability density (HPD) from these 500 datasets using the *HDInterval* v0.1.3 package (Meredith and Kruschke 2016) in R.

### ddRAD-Seq Demographic Modeling

While our WGS had many more loci, there was limited geographic representation, as well as fewer individuals sampled; therefore, we built regional models from the ddRAD-Seq dataset to explore additional population tree topologies. We estimated the SFS of each population in ANGSD using the reference aligned Illumina reads instead of the previously called SNPs.

We built regional models within the evolutionary clusters for eastern Asia/Pacific, SE Asia, Northern Europe, and Western Europe. We used this reductive approach to limit the number of parameters being estimated. Within each region, we compared topologies between populations suggested by previous population structure analyses (Puckett et al. 2016). We used the same fastsimcoal2 run parameters as described above; however, we did not create pseudo-observed datasets for parameter estimation, unless noted, as our interest was in topology. A secondary reason we did not further explore population parameters within the regions was that we observed these datasets tended to overestimate divergence times, likely due to unsorted variation remaining within populations until coalescence with the unsampled ancestral population. Finally, we investigated population topology and admixture proportions in Vancouver, Canada and Berkeley, USA since each site was identified as admixed in our previous analysis (Table S2).

### Chromosomal Diversity

Using the genotypes from the WGS data created with ANGSD, we estimated heterozygosity on each chromosome for each individual. We exported the genotype likelihoods into PLINK v1.9 (Purcell et al. 2007; Chang et al. 2015) and estimated heterozygosity (--het) on each chromosome.

## DATA ACCESS

Data for whole genome sequences from 10 brown rats available on NCBI SRA BioProject PRJNA344413.

## ACKNOWLEDGEMENTS

We thank Joshua Schraiber and two anonymous reviewers for comments that improved the manuscript. This work was funded by National Science Foundation grants DEB 1457523 and MRI 1531639 to JM-S. The mammal collections at the University of Alaska Museum of the North, University of California-Berkeley Museum of Vertebrate Zoology, the Burke Museum at the University of Washington, and the Museum of Texas Tech University also graciously provided tissue samples.

## REFERENCES

Aplin KP, Suzuki H, Chinen AA, Chesser RT, ten Have J, Donnellan SC, Austin J, Frost A, Gonzalez JP, Herbreteau V et al. 2011. Multiple geographic origins of commensalism and complex dispersal history of black rats. PLoS ONE 6: e26357.

Armitage P. 1993. Commensal rats in the New World, 1492-1992. Biologist 40: 174–178.

Atanur Santosh S, Diaz Ana G, Maratou K, Sarkis A, Rotival M, Game L, Tschannen Michael R, Kaisaki Pamela J, Otto Georg W, Ma Man Chun J et al. 2013. Genome Sequencing Reveals Loci under Artificial Selection that Underlie Disease Phenotypes in the Laboratory Rat. Cell 154: 691–703.

Beerli P. 2004. Effect of unsampled populations on the estimation of population sizes and migration rates between sampled populations. Molecular Ecology 13: 827–836.

Black L. 1983. Record of maritime disasters in Russian America, Part One: 1741-1799. In Proceedings of the Alaska Maritime Archaeology Workshop, May 17-19, 1983, Vol Alaska Sea Grant Report No. 83–9. University of Alaska-Fairbanks, Sitka, AK.

Chang CC, Chow CC, Tellier LCAM, Vattikuti S, Purcell SM, Lee JJ. 2015. Second-generation PLINK: rising to the challenge of larger and richer datasets. GigaScience 4: 7.

Combs M, Puckett EE, Richardson J, Mims D, Munshi-South J. 2018. Spatial population genomics of the brown rat (Rattus norvegicus) in New York City. Molecular Ecology 27: 83–98.

Deinum EE, Halligan DL, Ness RW, Zhang Y-H, Cong L, Zhang J-X, Keightley PD. 2015. Recent evolution in Rattus norvegicus is shaped by declining effective population size. Molecular Biology and Evolution 32: 2547–2558.

Ervynck A. 2002. Sedentism or urbanism? On the origin of the commensal black rat (Rattus rattus). In Bones and the man: Studies in honour of Don Brothwell, (ed. K Dobney, T O’Connor), pp. 95–109. Oxbow Books, Oxford.

Gibbs RA Weinstock GM Metzker ML Muzny DM Sodergren EJ Scherer S Scott G Steffen D Worley KC Burch PE et al. 2004. Genome sequence of the Brown Norway rat yields insights into mammalian evolution. Nature 428: 493–521.

Harper GA, Bunbury N. 2015. Invasive rats on tropical islands: Their population biology and impacts on native species. Global Ecology and Conservation 3: 607–627.

Heng D. 2009. Sino-Malay Trade and Diplomacy from the Tenth through the Fourteenth Century. Ohio University Press, Athens, USA.

Himsworth CG, Parsons KL, Jardine C, Patrick DM. 2013. Rats, cities, people, and pathogens: a systematic review and narrative synthesis of literature regarding the ecology of rat-associated zoonoses in urban centers. Vector borne and zoonotic diseases (Larchmont, NY) 13: 349–359.

Johnson MTJ, Munshi-South J. 2017. Evolution of life in urban environments. Science 358.

Jones HP, Holmes ND, Butchart SHM, Tershy BR, Kappes PJ, Corkery I, Aguirre-Muñoz A, Armstrong DP, Bonnaud E, Burbidge AA et al. 2016. Invasive mammal eradication on islands results in substantial conservation gains. Proceedings of the National Academy of Sciences 113: 4033–4038.

Korneliussen TS, Albrechtsen A, Nielsen R. 2014. ANGSD: Analysis of Next Generation Sequencing Data. BMC Bioinformatics 15: 1–13.

Lack J, Hamilton M, Braun J, Mares M, Van Den Bussche R. 2013. Comparative phylogeography of invasive Rattus rattus and Rattus norvegicus in the U.S. reveals distinct colonization histories and dispersal. Biological Invasions 15: 1067–1087.

Langmead B, Salzberg SL. 2012. Fast gapped-read alignment with Bowtie 2. Nature Methods 9: 357–359.

Li H. 2009. SNPable Regions. http://lh3lh3.users.sourceforge.net/snpable.shtml.

Li H, Durbin R. 2010. Fast and accurate long-read alignment with Burrows-Wheeler transform. Bioinformatics 26: 589–595.

Li H, Handsaker B, Wysoker A, Fennell T, Ruan J, Homer N, Marth G, Abecasis G, Durbin R. 2009. The Sequence Alignment/Map format and SAMtools. Bioinformatics 25: 2078–2079.

Lieberman V. 2009. Strange Parallels: Southeast Asia in Global Context, c. 800-1830. Cambridge University Press, Cambridge, Great Britain.

McKenna A, Hanna M, Banks E, Sivachenko A, Cibulskis K, Kernytsky A, Garimella K, Altshuler D, Gabriel S, Daly M et al. 2010. The Genome Analysis Toolkit: A MapReduce framework for analyzing next-generation DNA sequencing data. Genome Research 20: 1297–1303.

Meredith M, Kruschke J. 2016. Highest (Posterior) Density Intervals. CRAN, CRAN.

Moreno-Mayar JV, Potter BA, Vinner L, Steinrücken M, Rasmussen S, Terhorst J, Kamm JA, Albrechtsen A, Malaspinas A-S, Sikora M et al. 2018. Terminal Pleistocene Alaskan genome reveals first founding population of Native Americans. Nature 553: 203.

Peter BM, Slatkin M. 2013. Detecting range expansions from genetic data. Evolution 67: 3274–3289.

Pimentel D, Lach L, Zuniga R, Morrison D. 2000. Environmental and Economic Costs of Nonindigenous Species in the United States. BioScience 50: 53–65.

Puckett EE, Micci-Smith O, Munshi-South J. 2018. Genomic analyses identify multiple Asian origins and deeply diverged mitochondrial clades in inbred brown rats (Rattus norvegicus). Evolutionary Applications 11: 718–726.

Puckett EE, Park J, Combs M, Blum MJ, Bryant JE, Caccone A, Costa F, Deinum EE, Esther A, Himsworth CG et al. 2016. Global population divergence and admixture of the brown rat (Rattus norvegicus). Proceedings of the Royal Society B: Biological Sciences 283: 1–9.

Purcell S, Neale B, Todd-Brown K, Thomas L, Ferreira MAR, Bender D, Maller J, Sklar P, de Bakker PIW, Daly MJ et al. 2007. PLINK: A tool set for whole-genome association and population-based linkage analyses. The American Journal of Human Genetics 81: 559–575.

Scheet P, Stephens M. 2006. A fast and flexible statistical model for large-scale population genotype data: Applications to inferring missing genotypes and haplotypic phase. American Journal of Human Genetics 78: 629–644.

Schiffels S, Durbin R. 2014. Inferring human population size and separation history from multiple genome sequences. Nature Genetics 46: 919–925.

Slatkin M. 2005. Seeing ghosts: the effect of unsampled populations on migration rates estimated for sampled populations. Molecular Ecology 14: 67–73.

Smith AT, Xie Y. 2008. A Guide to the Mammals of China. Princeton University Press, Princeton, NJ.

Song Y, Lan Z, Kohn MH. 2014. Mitochondrial DNA phylogeography of the Norway rat. PLoS ONE 9: e88425.

Suzuki H, Nunome M, Kinoshita G, Aplin KP, Vogel P, Kryukov AP, Jin ML, Han SH, Maryanto I, Tsuchiya K et al. 2013. Evolutionary and dispersal history of Eurasian house mice Mus musculus clarified by more extensive geographic sampling of mitochondrial DNA. Heredity 111: 375–390.

Tucker J. 2015. The Silk Road: China and the Karakorum Highway. I.B.Tauris & Company, New York, USA.

Yalden DW. 2003. Mammals in Britain – A historical perspective. British Wildlife 14: 243–251.

Zeng L, Ming C, Li Y, Su L-Y, Su Y-H, Otecko NO, Dalecky A, Donnellan S, Aplin K, Liu X-H et al. 2018. Out of Southern East Asia of the Brown Rat Revealed by Large-Scale Genome Sequencing. Molecular Biology and Evolution doi:10.1093/molbev/msx276.

